# MetaLigand: A database for predicting non-peptide ligand mediated cell-cell communication

**DOI:** 10.1101/2025.01.14.633094

**Authors:** Ying Xin, Yang Jin, Cheng Qian, Seth Blackshaw, Jiang Qian

## Abstract

Non-peptide ligands (NPLs), including lipids, amino acids, carbohydrates, and non-peptide neurotransmitters and hormones, play a critical role in ligand-receptor-mediated cell-cell communication, driving diverse physiological and pathological processes. To facilitate the study of NPL-dependent intercellular interactions, we introduce MetaLigand, an R-based and web-accessible tool designed to infer NPL production and predict NPL-receptor interactions using transcriptomic data. MetaLigand compiles data for 233 NPLs, including their biosynthetic enzymes, transporter genes, and receptor genes, through a combination of automated pipelines and manual curation from comprehensive databases. The tool integrates both de novo and salvage synthesis pathways, incorporating multiple biosynthetic steps and transport mechanisms to improve prediction accuracy. Comparisons with existing tools demonstrate MetaLigand’s superior ability to account for complex biogenesis pathways and model NPL abundance across diverse tissues and cell types. Furthermore, analysis of single-nucleus RNA-seq datasets from age-related macular degeneration samples revealed that distinct retinal cell types exhibit unique NPL profiles and participate in specific NPL-mediated pathological cell-cell interactions. Finally, MetaLigand supports single-cell RNA sequencing (scRNA-seq) and spatial transcriptomics data, enabling the visualization of predicted NPL production levels and heterogeneity at single-cell resolution.

## INTRODUCTION

Cell-cell communication (CCC) is fundamental to the organization and functioning of multicellular organisms, enabling cells to coordinate activities for biological processes and homeostasis^1^. A primary mechanism of CCC is ligand-receptor interaction (LRI), where a ligand binds its specific receptor to trigger intracellular signaling cascades, modulate cellular actions, and regulate gene expression. The complexity of such interactions underpins the intricate functional machinery of metazoans, demanding systematic exploration for deeper understanding.

Advances in computational pipelines and databases have enhanced the modeling of CCC, enabling the profiling of its quantitative, dynamic, and heterogeneous aspects in tissues. Single-cell sequencing technologies now allow unbiased, simultaneous sequencing of cells within tissues, offering a comprehensive view of transcriptomic landscapes^2^. Single-cell transcriptomic analyses have proven particularly useful for investigating LRIs, using gene expression levels of ligands and receptors (LRs) as proxies for their abundance^3,4^. Tools such as NicheNet^5^, CellPhoneDB^6^, CellChat^7^, and LRLoop^8,9^ have greatly advanced our understanding of LRI roles in cellular physiology, development, and disease progression.

However, these methods primarily focus on peptide ligands encoded by single genes, neglecting non-peptide ligands (NPLs), molecules like non-peptide hormones, neurotransmitters, amino acids, lipids, and carbohydrates. Since NPLs are not genome-encoded or transcribed into mRNA, they are difficult to analyze directly from transcriptomic data and have been largely overlooked in computational studies. The production of small-molecule ligands involves sequential enzymatic reactions transforming precursor molecules, with additional steps such as transporter-mediated shuttling or vesicular release for hydrophilic ligands^10^. Modeling NPL production based on the expression of genes involved in these processes, such as synthetic enzymes, and transporters, opens the possibility of transcriptomic-based analyses of NPL-mediated CCC.

Previous attempts to model NPL-based LRIs include NeuronChat^11^, which focused on eight neurotransmitters, using manually curated gene sets of enzymes and transporters. Tools like CellPhoneDBv5^12^ and MRCLinkdb^10^ have expanded this concept by curating databases for over 100 NPLs, incorporated into pipelines such as CellChatv2^13^. However, these methods have limitations. They often rely on manual curation or databases like HMDB, which lack detailed metabolic pathway annotations^14^. Furthermore, they overlook alternative ligand production routes, such as salvage pathways (e.g., dopamine synthesis via L-DOPA uptake^15^, synthesis of the amino acid ligands such as L-Arginine and Ornithine from the influx transported precursors^16^). Finally, existing models have not been extensively validated for NPL abundance predictions across tissues and cell types.

In this study, we present MetaLigand, an R-based, customizable bioinformatics tool for profiling CCC via NPL-receptor interactions using transcriptomic data. MetaLigand compiles data for 233 NPLs, integrating gene sets for biosynthetic enzymes, transporters, and receptors. Biogenesis pathway information is extracted from genome-scale metabolic models (GEMs) in the Metabolic Atlas database^17,18^, followed by manual curation. MetaLigand’s predictions of NPL production can be supported by literatures and mass spectrometry datasets. The tool enables users to infer, visualize, and analyze NPL-dependent LRIs across bulk RNA-seq, single-cell RNA-seq (scRNA-seq), and spatial transcriptomic data.

## RESULTS

### Data Collection Pipeline

MetaLigand integrates data through an automated pipeline combined with manual curation (Figure 1). The database compiles non-peptide ligand (NPL)-receptor interactions across human, mouse, and zebrafish species. Small molecule-protein interactions are sourced from databases such as BindingDB^19^, Cellinker^20^, BioGRID^21^, STITCH^22^, CTD^23^, and CellphoneDBv5^12^. Endogenous metabolite data are retrieved from the HMDB database^14^. Receptor genes are obtained from OmniPath^24^, CellTalkDB^25^, Cellinker^20^, Cell-Cell Interaction Database^26^, NicheNet^5^, and NATMI^27^, with cross-referencing against GeneCards^28^ and HGNC^29^ databases.

**Fig. 1.**
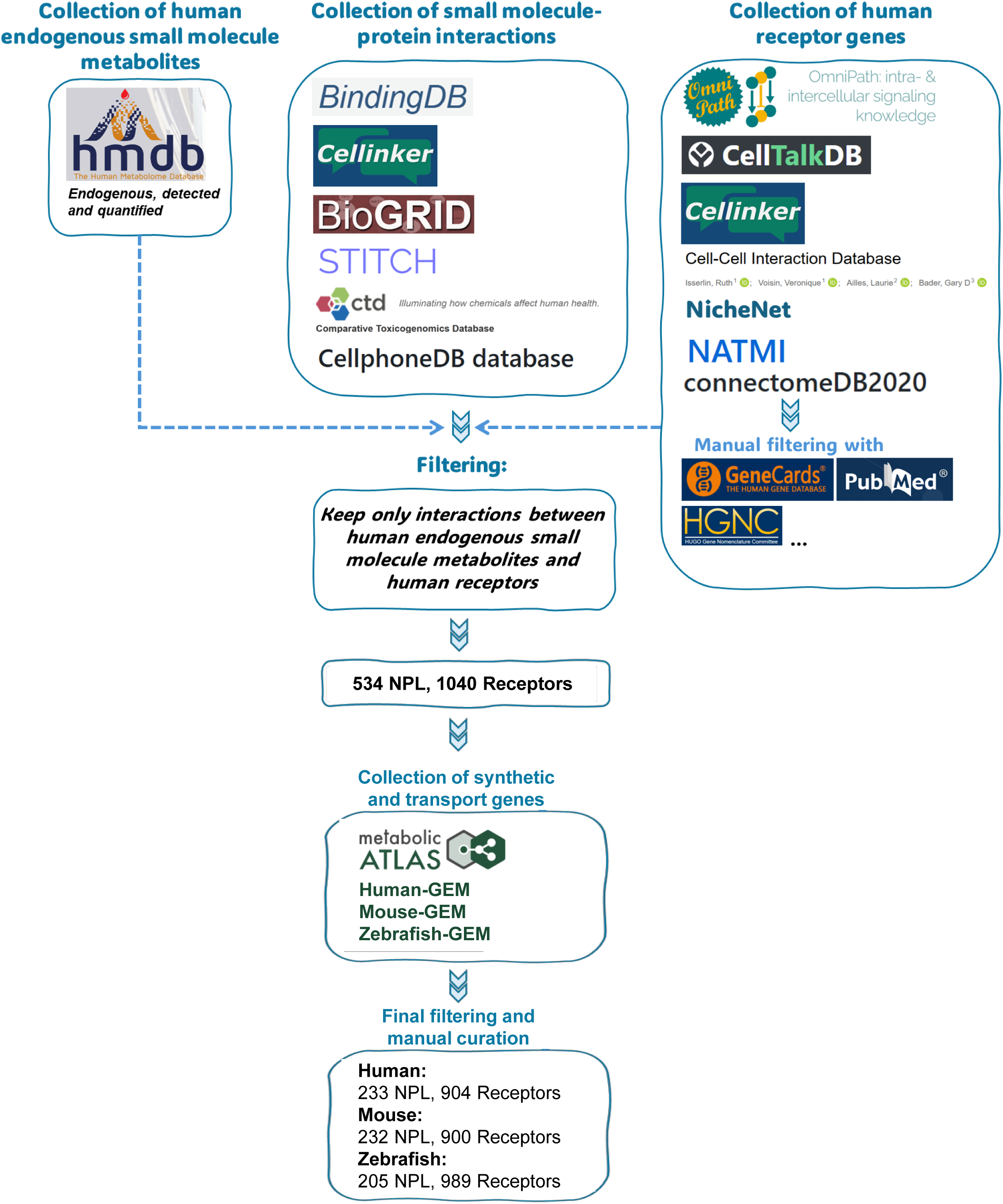
Curation of the MetaLigand database. MetaLigand integrates data through an automated pipeline combined with manual curation. Small molecule-protein interactions are sourced from databases including BindingDB, Cellinker, BioGRID, STITCH, CTD, and CellphoneDBv5. Endogenous metabolite data are retrieved from the HMDB database. Receptor genes are obtained from OmniPath, CellTalkDB, Cellinker, Cell-Cell Interaction Database, NicheNet, and NATMI, with cross-referencing against GeneCards and HGNC databases. Small molecule-protein interactions are then filtered to include only those between human endogenous metabolite NPLs and their receptors, resulting in 13,441 interaction pairs involving 534 NPLs and 1,040 receptors. These 534 NPLs are further queried in the GEM of Metabolic Atlas database for relevant metabolic pathway reactions and enzymes with predefined criteria to identify those involved in metabolite synthesis, secretion, and transport. This auto-generated NPL database is then manually curated further using literature and databases such as TCDB. After excluding NPLs that cannot be endogenously synthesized or are challenging to estimate from transcriptomic data, MetaLigand includes 233 NPLs for humans, 232 for mice, and 205 for zebrafish, as well as 904, 900, and 989 receptors that interact with the NPLs for humans, mice, and zebrafish, respectively.

Small molecule-protein interactions are filtered to include only those between human endogenous metabolite NPLs and their corresponding receptors, resulting in 13,441 interaction pairs involving 534 NPLs and 1,040 receptors (Figure 1). These 534 NPLs are further processed through an automated pipeline to identify synthetic enzymes and membrane transporters involved in their biogenesis and bioavailability. Synonyms for each NPL are gathered from HMDB^14^, and corresponding GEM names are used to query the Metabolic Atlas database^17^ for relevant metabolic pathway reactions and enzymes. The reactions are filtered using predefined criteria to identify those involved in metabolite synthesis, secretion, and transport. This auto-generated NPL database is curated further using literature and databases such as TCDB^30^.

After excluding NPLs that cannot be endogenously synthesized (e.g., linolenic acid and tryptophan) or are challenging to estimate from transcriptomic data (e.g., water), MetaLigand includes 233 NPLs for humans, 232 for mice, and 205 for zebrafish (Figure 1). It also contains 904 receptors for humans, 900 for mice, and 989 for zebrafish, that interact with the NPLs (Figure 1).

### Overview of MetaLigand

In the MetaLigand database, approximately 60% of NPLs are associated with synthetic enzymes, while the remaining NPLs are linked to both synthetic enzymes and membrane transporters (Figure 2A). Biochemically, most NPLs included in MetaLigand are lipids, amino acids, and nucleic acids, with smaller but functionally significant groups represented by non-peptide hormones and neurotransmitters (Figure 2B). Each NPL is associated with one or more genes encoding five key protein groups: (1) enzymes for NPL synthesis (Synthetic step 1), (2) enzymes for synthesizing NPL precursors (Synthetic step 2), (3) transporters for secreting NPLs towards the extracellular space and/or loading the synaptic vesicles (Efflux step), (4) uptake transporters for importing NPL precursors from extracellular spaces (Influx step), and (5) NPL receptors (Figure 2C).

**Fig. 2.**
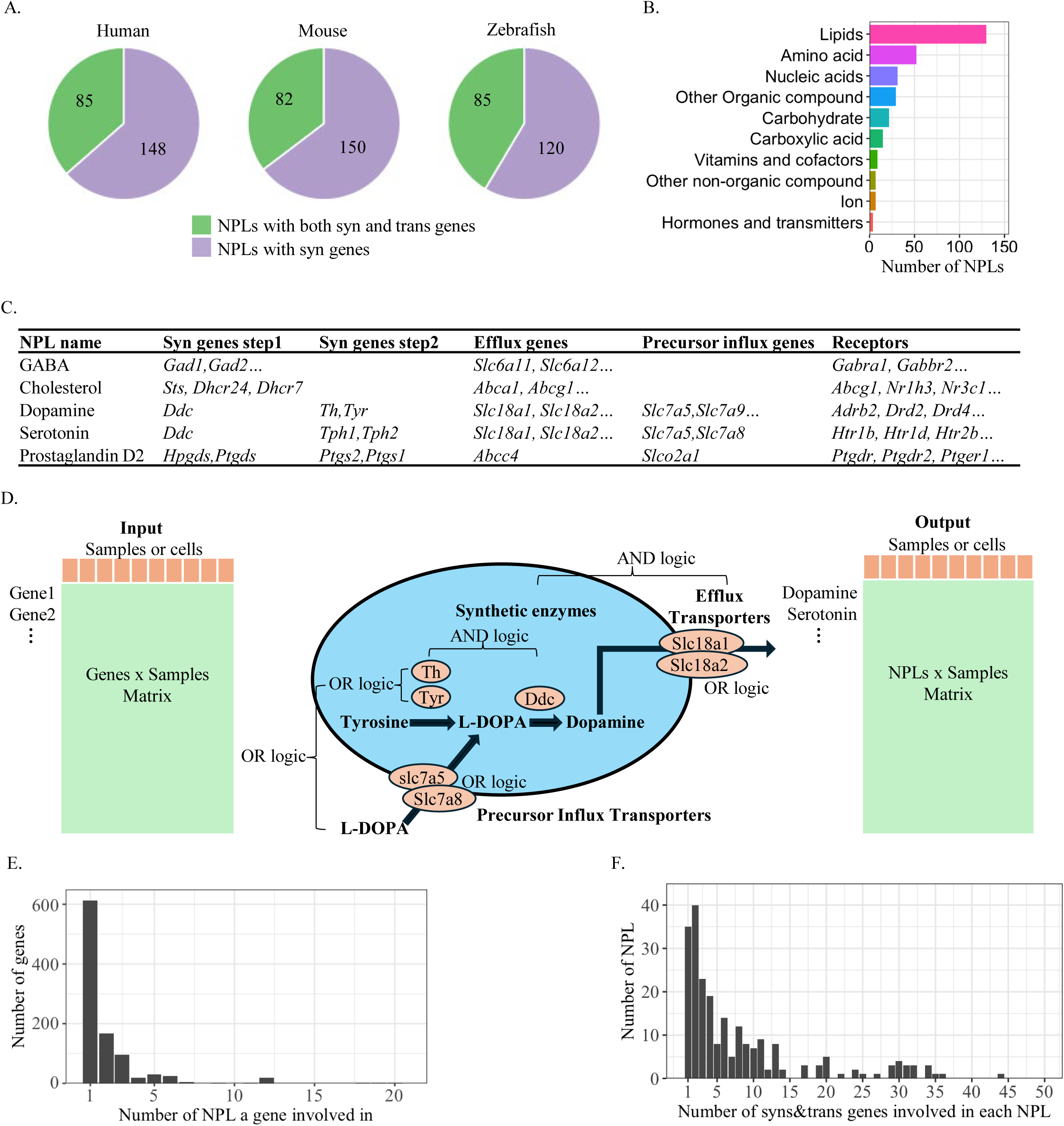
Overview of MetaLigand. **(A)** Composition of NPLs in the MetaLigand database in terms of their association with synthetic enzymes and transporters. In the MetaLigand database, approximately 59%∼65% of NPLs are associated with synthetic enzymes for human, mouse, and zebrafish species, while the remaining NPLs are linked to both synthetic enzymes and membrane transporters. **(B)** Biochemical categories of the NPLs curated in the MetaLigand database. Most of the NPLs in MetaLigand are lipids, amino acids, and nucleic acids, other smaller yet functionally significant groups include non-peptide hormones, neurotransmitters, etc. **(C)** Structure of the MetaLigand database. Each NPL is associated with genes encoding five protein groups: (1) enzymes for NPL synthesis (Syn genes step 1), (2) enzymes for synthesizing NPL precursors (Syn genes step 2), (3) transporters for secreting NPLs towards the extracellular space and/or loading the synaptic vesicles (Efflux genes), (4) uptake transporters for importing NPL precursors from extracellular spaces (Precursor influx genes), and (5) NPL receptors **(D)** Schematic diagram of the computational model of MetaLigand for the inference of NPL production from transcriptomic data. The abundance of each NPL is modeled based on the expression levels of the genes in the first four groups. Within each group, gene expressions are summarized using “OR logic” (arithmetic mean); NPL synthesis predictions consider the final two steps of the synthesis pathway (Syn step 1 and Syn step 2), linked by “AND logic” (geometric mean); Salvage synthesis pathways, where cells recycle or import NPL precursors from the extracellular space (Precursor influx) and process them into final NPL products using last-step enzymes (Syn step 1), are also included, connected with the de novo pathways by “OR logic”; Efflux transporters (Efflux genes), when required for the secretion of synthesized NPLs, are linked to synthesis pathways using “AND logic”; NPLs without associated transporters in referenced databases are assumed to be directly released into the extracellular space after biosynthesis. **(E-F)** Histograms exhibit specificity of the association between NPLs and the relevant gene sets in MetaLigand mouse model. (E) Distribution of the number of NPLs the genes involved in. (F) Distribution of the number of synthetic enzymes and transporter genes involved in each NPL.

The abundance of each NPL is modeled based on the expression levels of the genes in the first four groups (Figure 2D). (1) Within each group, gene expression is summarized using “OR logic” (arithmetic mean). (2) NPL synthesis predictions consider the final two steps of the synthesis pathway (Synthetic step 1 and Synthetic step 2), which are linked by “AND logic” (geometric mean), ensuring that sequential catalytic reactions must both be active to produce the final NPL. (3) Salvage synthesis pathways, where cells recycle or import NPL precursors from the extracellular space (Influx step) and process them into final NPL products using last-step enzymes (Synthetic step 1), are also included. The de novo and salvage pathways are connected by “OR logic.” (4) The secretion of synthesized NPLs often requires efflux transporters (Efflux step), which are linked to synthesis pathways using “AND logic.” For NPLs not associated with transporters in referenced databases, it is assumed that these NPLs are directly released into the extracellular space after biosynthesis.

Regarding the specificity of association between NPLs and their relevant gene sets (synthetic enzymes, transporters, etc.), an example from the mouse model shows that 618 (61.1%) of the 1012 enzyme and transporter genes in MetaLigand are uniquely associated with the production of a single NPL (Figure 2E). In total, 125 NPLs (53.9%) are linked to fewer than five genes, while 195 NPLs (84.1%) are represented by fewer than 15 genes (Figure 2F). Similar trends are observed in human and zebrafish models (Suppl. Figure 1).

### A Non-Peptide Ligand Atlas for Mouse

Using MetaLigand, we analyzed a public single-cell sequencing dataset including 975 cell types in 36 tissues^31–33^. This analysis generated a single-cell resolution NPL atlas for the mouse. As anticipated, nervous tissues such as the neocortex and retina exhibited distinct NPL patterns compared to other tissues (Figure 3A). Additionally, metabolic organs like the kidney, liver, and adipose tissue showed a broader diversity of NPLs, particularly when compared to structural tissues such as bone or muscle (Figure 3A)

**Fig. 3.**
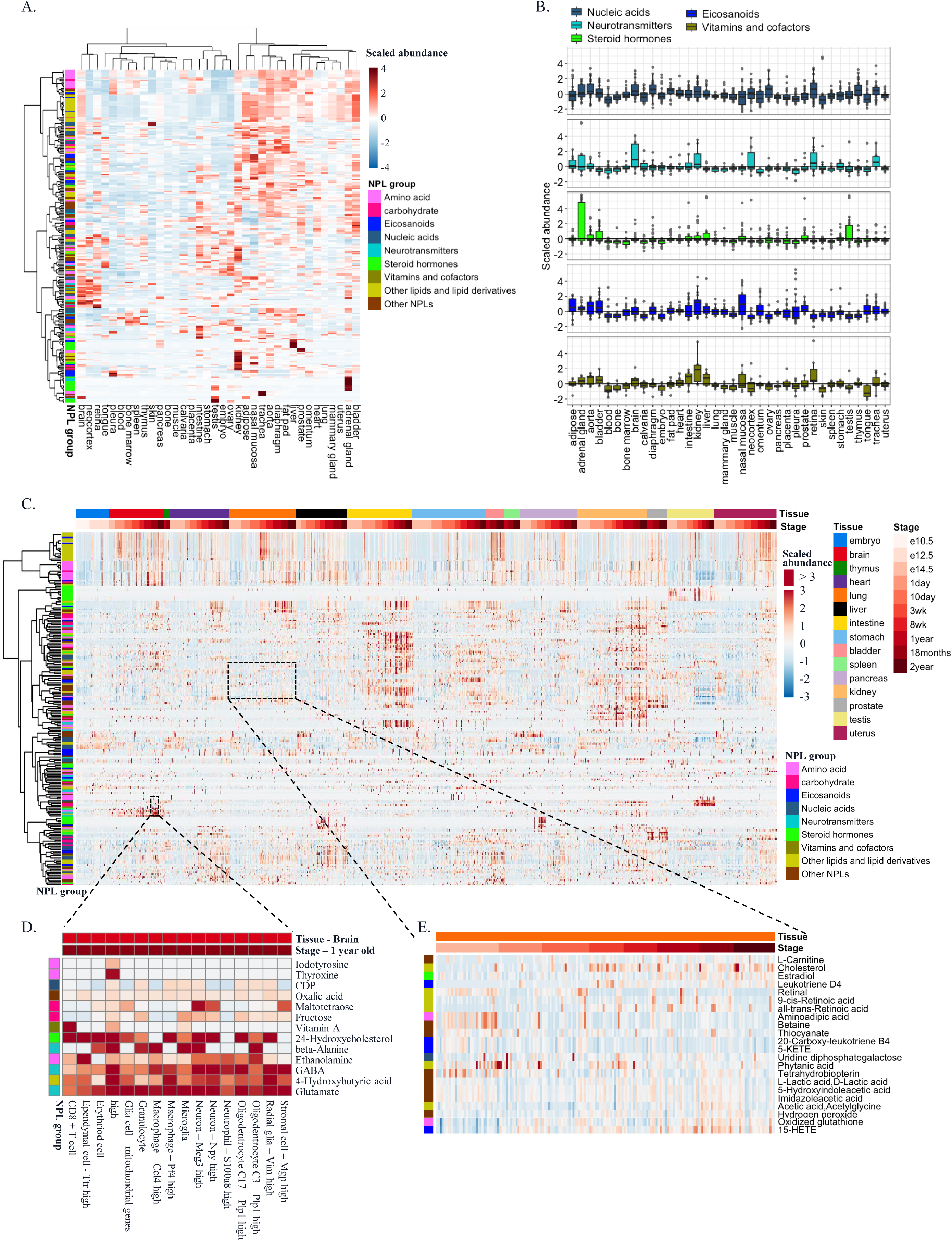
A Non-Peptide Ligand Atlas for Mouse generated by MetaLigand. **(A)** Heatmap exhibits NPL patterns across different organ tissues. **(B)** Patterns of the biosynthesis of various NPL categories, including nucleic acids, neurotransmitters, steroid hormones, eicosanoids, vitamins and cofactors, across different tissues. **(C)** Applying MetaLigand to the MCA v1.0, v2.0, and v3.0 datasets generates dynamics of various NPL categories across ten developmental stages in fifteen different tissues. **(D)** Heatmap shows sample NPL predictions in brain cell types in one-year-old stage. **(E)** Heatmap of sample NPL predictions in various lung cell types across multiple stages.

The biosynthesis of each NPL category demonstrated unique patterns across different tissues (Figure 3B). For instance, neurotransmitters are abundant in neuronal tissues such as the brain and retina, and a few peripheral organs such as the kidney; Vitamins are most prevalent in the kidney and retina; Steroid hormones accumulate primarily in the adrenal glands and testes; Eicosanoids are most abundant in the nasal mucosa, where they are produced during inflammatory responses. Many of these observations are supported by previous studies^34^ ^35^ ^36^.

We then examine the dynamics of NPL abundance across 10 developmental stages in 15 tissues using the MCA v1.0, v2.0, and v3.0 datasets ^32,33^ (Figure 3C). In addition to the high cell-type-specificity, such as the predicted high abundance of neurotransmitters like GABA and glutamate in brain-originated cell types (Figure 3D) and that leukotrienes and lipoxins are predominantly produced by immune cells including CD8+ T cells, macrophages, and microglia, many NPLs exhibited development stage-dependent patterns. For example, hydroxyeicosatetraenoic acids like 15-HETE, key pro-inflammatory mediators, showed increased levels across various cell types in aged lung tissue, consistent with previous findings^37^. Similarly, cholesterol derivatives also displayed elevated levels in aging lung tissue, aligning with the upregulation of cholesterol biosynthesis observed in aged lungs and its involvement in lung fibrosis^38^ (Figure 3C and 3E).

### Comparing MetaLigand to Other Methods

To evaluate the performance of MetaLigand, we compared it to three computational methods and their associated databases: MRClinkDB^10^, CellphoneDBv5^12^, and NeuronChat^11^. MetaLigand includes a significantly larger number of NPLs and integrates a more biologically meaningful model overall (Figure 4A and Figure 2D). While NeuronChat utilizes multiple synthetic enzyme genes, it is limited to only eight neurotransmitters. In contrast, MRClinkDB and CellphoneDBv5 cover a broader range of NPLs but model NPL production based only on a single step in the synthetic pathway (Figure 4B).

**Fig. 4.**
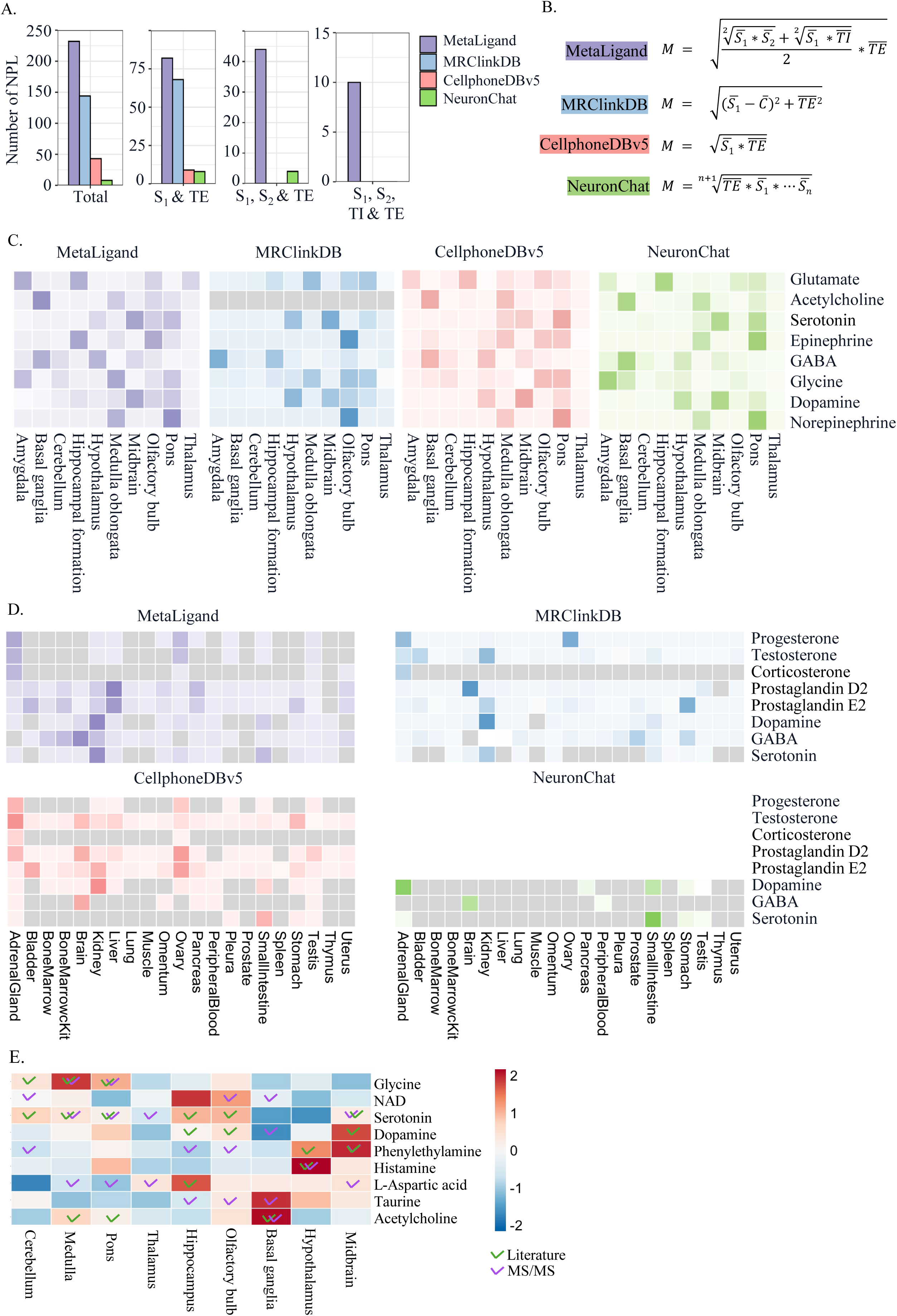
Comparison of MetaLigand to other methods in the prediction of NPL production. **(A)** An overview on the number of NPLs and their association with the synthetic and transporter gene groups in the databases of MetaLigand, MRClinkDB, CellphoneDBv5, and NeuronChat. **(B)** Computational models of NPL prediction through transcriptomic data used in MetaLigand, MRClinkDB, CellphoneDBv5, and NeuronChat. **(C)** Heatmaps exhibit predictions of the physiological distribution of neurotransmitters in nervous tissues using the mouse brain atlas by MetaLigand, MRClinkDB, CellphoneDBv5, and NeuronChat. **(D)** Heatmaps of sample NPL predictions in multiple mouse organs using the MAC1 gene expression atlas by MetaLigand, MRClinkDB, CellphoneDBv5, and NeuronChat. **(E)** Comparison of neurotransmitter NPL predictions by MetaLigand in different brain regions using scRNA-seq dataset from an adult mouse brain atlas, to literature reports and mass spectrometry (MS)-based metabolomics data obtained from the mouse brain.

First, we assessed MetaLigand’s capability to capture the physiological distribution and abundance of neurotransmitters in nervous tissues using the mouse brain atlas^39^ (Figure 4C). MetaLigand, CellphoneDBv5, and NeuronChat successfully detected all eight neurotransmitters across brain regions, while MRClinkDB failed to detect acetylcholine. Notably, MetaLigand accurately identified high glutamate abundance in the thalamus, a region enriched with excitatory glutamatergic neurons^40,41^. Furthermore, MetaLigand and MRClinkDB also successfully identify high glycine abundance in the medulla oblongata, a region rich in inhibitory glycinergic neurons^42^, whereas the other two methods failed to detect glycine in this region.

Beyond neurotransmitters, we compared predictions for steroid hormones, a particularly challenging category for transcriptomics-based modeling due to their ubiquitous biosynthetic enzymes facilitating diverse NPL production. We used the MAC1 gene expression atlas in mouse organs for the prediction ^43^ (Figure 4D). MetaLigand show progesterone, testosterone, and corticosterone present in adrenal glands, ovaries, and testis, all of which are the major steroidogenic tissues^44^. In contrast, CellphoneDBv5 predicts high testosterone in almost all organs, including the stomach which is a non-steroidogenic tissue. Similarly, MRClinkDB incorrectly predicts higher testosterone in the bladder and kidney. NeuronChat, being limited to neurotransmitters, was incapable of characterizing steroid hormones.

MetaLigand’s consideration of both de novo and salvage synthesis pathways further enhanced its predictive accuracy. For example, dopamine can be synthesized de novo in dopaminergic neurons or via salvage pathways in non-neuronal tissues such as the kidney and intestine^45^. MetaLigand addresses this by including two synthesis steps for *de novo* synthesis and influx transporter genes for salvage pathway. As a result, it identified high dopamine level in the brain, kidney, and intestine (Figure 4D). In comparison, NeuronChat, which does not account for salvage pathways, failed to profile dopamine production in the kidney and intestine. Similarly, the prostaglandin E2 (PGE2) and prostaglandin D2 (PGD2) can be synthesized from the uptake of the precursor prostaglandin H2 (PGH2), both of which are highly involved in inflammation, pain sensation and other biological processes in various tissues^46,47^. MetaLigand can detect PGE2 and PGD2 in a wide spectrum of organs such as liver, kidney, and pancreas, enabling comprehensive modeling of NPL abundance and dynamics (Figure 4D).

In addition to computational predictions, metabolite abundance is often quantified using mass spectrometry (MS)^48^. To validate MetaLigand’s NPL predictions, we applied MetaLigand to a scRNA-seq data from an adult mouse brain atlas^39^ and compared the predictions to MS-based metabolomics data obtained from the mouse brain^49^. Focusing on neurotransmitters, MetaLigand’s prediction showed strong agreement with MS measurements and literature reports (Figure 4E). For instance, MetaLigand identified high glycine abundance in the medulla and pons, consistent with MS data and previous studies showing glycinergic neurons in these regions^49^. However, MS sometimes fails to distinguish the synthetic origins and active sites of the NPLs. For example, MetaLigand predicted dopamine synthesis in the midbrain, while MS data indicates high abundance of dopamine in the basal ganglia (Figure 4E). This discrepancy likely arises because dopaminergic neurons, located in the substantia nigra pars compacta (SNc) and ventral tegmental area (VTA) of the midbrain^50,51^, release dopamine to target regions in the basal ganglia, where it regulates motor behavior^52^. This example highlights MS’s limitations as a gold standard for NPL prediction.

### Identifying Differential NPL-Mediated LRIs in Age-Related Macular Degeneration

Finally, we used MetaLigand to study the NPL-mediated cell-cell interactions in age-related macular degeneration (AMD), which is a progressive neurodegenerative disease and a leading cause of blindness worldwide^53^. To sustain constant visual signal transduction, photoreceptors require active metabolism to meet high energy demands and continuously synthesize their outer segment structures. Retinal pigment epithelium (RPE) cells, along with glial cells such as Müller cells, play essential roles in maintaining this homeostasis by supplying nutrients and clearing waste. Consequently, a more comprehensive characterization, involving diverse metabolite types, multiple cell types, pathological temporal dynamics, and intercellular communications, is essential for uncovering finer mechanisms of AMD^54^.

We applied MetaLigand to a public single-nucleus RNA-seq dataset comprising 164,399 cells from the retina, RPE, and choroid of seven control and six AMD donor eyes^55^ (Figure 5A). The analysis of NPLs across major cell types revealed the number of NPLs produced by each cell type (Suppl. Figure 2A). Notably, different cell types exhibited distinct panels of NPLs, suggesting that unique sets of small molecules may serve as ligands for intercellular communication (Figure 5B).

**Fig. 5.**
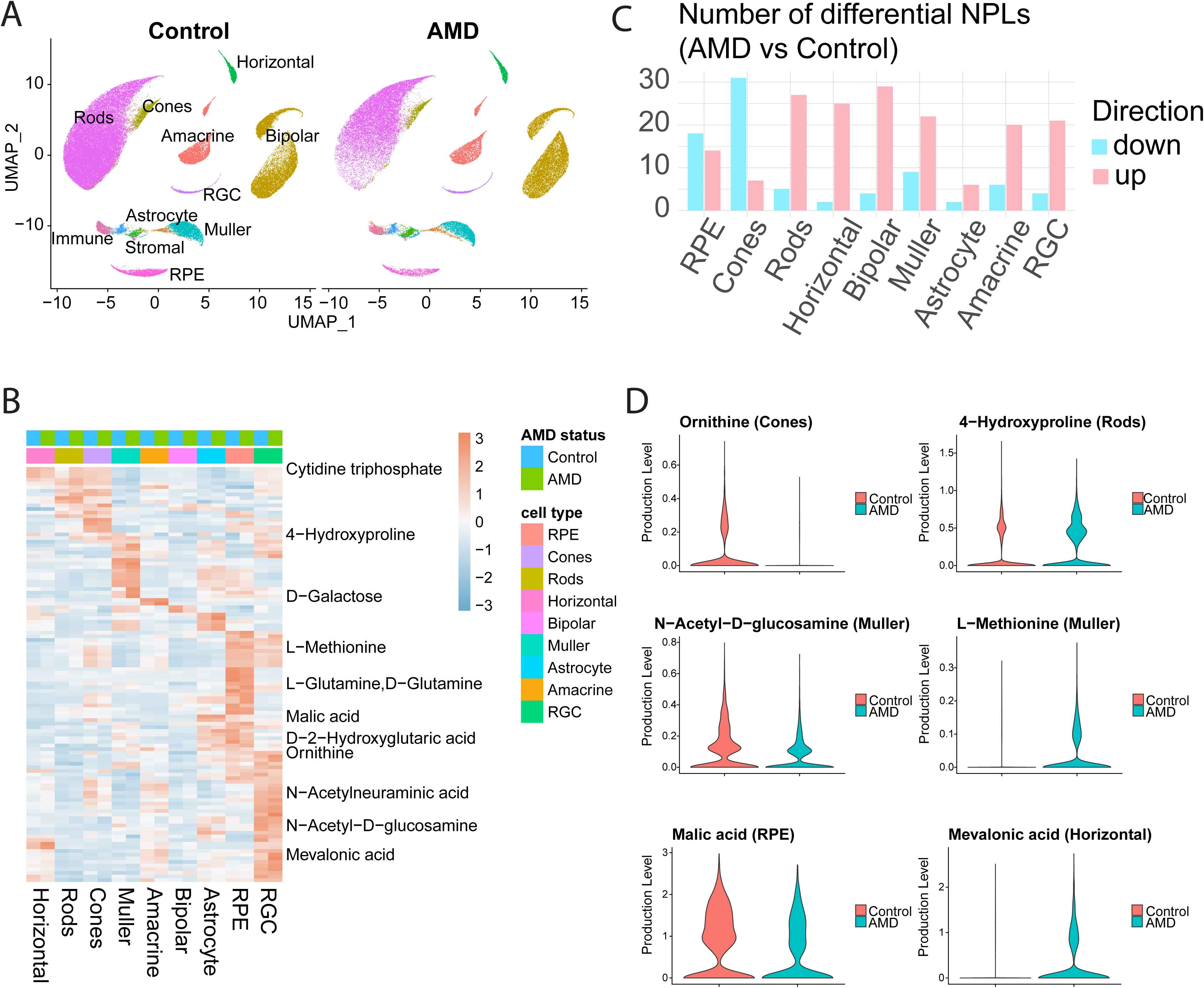
MetaLigand identifies differential NPLs in human retina AMD. **(A)** UMAP dimensionality reductions show cell types identified in a public single-nucleus RNA-seq dataset comprising 164,399 cells from the retina, RPE, and choroid of control and age-related macular degeneration (AMD) donor eyes. **(B)** Heatmap exhibits distinct panels of differential NPLs (detected in at least 10% of the cells in either of the clusters under comparison, with the absolute value of the average log2 fold change greater than 0.2 and the adjusted p-value from the Wilcoxon Rank Sum test below 0.05) between AMD and control in major retinal cell types. **(C)** Number of up- and down-regulated NPLs in AMD compared to control in major retinal cell types, colored in pink and blue, respectively. **(D)** Violin plots of predicted levels of sample identified differential NPLs in AMD and control cells in different cell types.

When comparing AMD to controls, we identified differential NPL regulation between AMD and controls. RPE cells and cones exhibited a greater number of downregulated than upregulated NPLs in AMD. In contrast, rod photoreceptors, various interneurons, Müller glia cells, and retinal ganglion cells (RGCs) showed increased NPL production in AMD compared to controls (Figure 5C). This observation suggests a potential compensatory homeostatic response by these cells to mitigate vision loss during AMD progression. Several of the identified differential NPLs are associated with AMD pathology. For example, ornithine was downregulated in cones and has been implicated in RPE and photoreceptor survival in AMD^56,57^. Similarly, glucosamine, which was downregulated in Müller cells, is considered a nutrient with potential protective effects against AMD^58–60^. In contrast, methionine, which was upregulated in Müller cells, has been suggested as a potential risk factor for AMD development^61–63^ (Figure 5D).

We identified differential NPL-mediated ligand-receptor interactions (LRIs) between AMD and controls, among six of the major cell types that make direct contacts with one another in adult retina. Figure 6A illustrates the predicted level of NPLs in sender cells and the gene expression levels of corresponding receptors in receiver cells. As expected, differential NPLs such as glucosamines, 4-hydroxyproline, L-methionine, and mevalonic acid are implicated in these differential ligand-receptor interactions. To further explore these interactions, we calculated the total number of differential NPL-mediated LRIs between retinal cell types (Figure 6B, 6C). Cone cells displayed the highest number of down-regulated LRIs among all cell types in AMD (Figure 6C) and the lowest number of up-regulated LRIs (Figure 6B). Notably, major cell types that lost connections with cone cells under AMD conditions include RPEs, bipolar cells, and Müller glia cells (Figure 6C). A comprehensive summary of cell-type-pair-specific LRIs is provided in Suppl. Figure 2B. Furthermore, Figures 6D and 6E highlight significant up- and down-regulated LRIs specific to certain cell types. Up-regulated, highly cell-type-pair-specific LRIs predominantly originate from Müller cells, RPEs, and rods (Figure 6D). Conversely, down-regulated, highly specific LRIs are most frequently observed between RPEs, cone cells, horizontal cells, and Müller cells (Figure 6E).

**Fig. 6.**
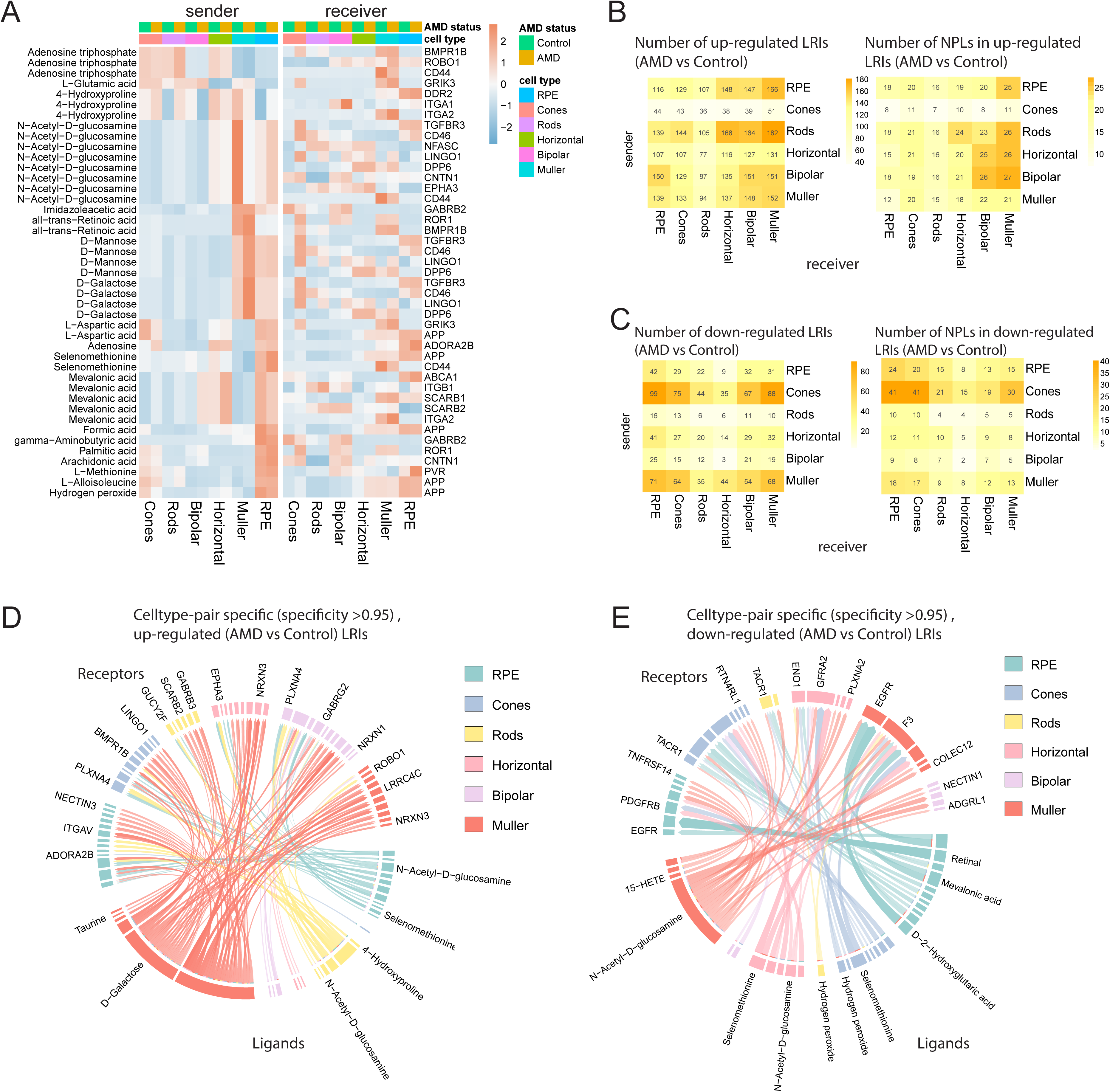
MetaLigand identifies differential NPL-mediated cell-cell interactions in human retina AMD. **(A)** Predicted NPL levels in sender cells and gene expression levels of the corresponding receptors in receiver cells of sample significant differential NPL-mediated ligand-receptor interactions (LRIs) (either the NPL or receptor gene detected in at least 25% of the cells in either of the clusters under comparison, with the absolute value of the average log2 fold change greater than 0.5 and the adjusted p-value from the Wilcoxon Rank Sum test below 0.01, while the interacting counterpart detected in at least 10% of the cells in the corresponding cluster and not differentially regulated in the opposite direction) between AMD and control in each cell type, each condition. **(B-C)** Number of up- and down-regulated NPL-mediated LRIs (either the NPL or receptor gene detected in at least 10% of the cells in either of the clusters under comparison, with the average log2 fold change greater than 0.2 or less than −0.2, respectively, and the adjusted p-value from the Wilcoxon Rank Sum test below 0.05, while the interacting counterpart detected in at least 10% of the cells in the corresponding cluster and not differentially regulated in the opposite direction) in AMD compared to control between each pair of sender and receiver cell types (left panels); Number of NPLs in the up- and down-regulated LRIs in AMD compared to control between each pair of cell types (right panels). **(D-E)** Significantly up- and down- regulated, highly cell-type-pair specific (with specificity score--the proportion of all cell-type pairs with the interaction scores of specified LR pair below the interaction score of the LR pair between the cell-type pair of focus--greater than 0.95) NPL-mediated LRIs.

## DISCUSSION

MetaLigand is a computational tool designed to infer non-peptide ligand (NPL) production and NPL-receptor interactions from transcriptomic data, including bulk RNA-seq, scRNA-seq, and spatial transcriptomics. By curating an extensive list of NPLs and their receptors, MetaLigand effectively addresses challenges in modeling NPL abundance using gene expression data. Its integration of synthetic enzymes and transport proteins into the predictive model makes it highly robust for analyzing complex cellular processes. Additionally, MetaLigand is compatible with popular computational tools for single-cell sequencing and spatial transcriptomics (Suppl. Figure 3). The full package, including NPL estimation and NPL-associated cell-cell communication (CCC) models, is available as an R package at https://github.com/jinyangye119/MetaLigand.

When compared to existing methods like NeuronChat^11^, MRClinkDB^10^, and CellphoneDBv5^12^, MetaLigand offers several key advantages. First, it incorporates both de novo and salvage synthetic pathways, providing comprehensive insights into NPL biosynthesis across cell types and tissues. Second, MetaLigand uses the geometric mean rather than the Euclidean norm to summarize genes involved in synthetic and transporter pathways, enhancing the biological relevance of its model. For ligands like neurotransmitters and steroid hormones, this approach captures the coordinated expression of synthetic enzymes and transporters, addressing disruptions that can lead to ligand accumulation or deficiency. In contrast, tools like MRClinkDB compute NPL abundance by subtracting catalytic enzyme expression from synthetic enzyme expression, which can result in inaccuracies. For instance, MRClinkDB failed to detect acetylcholine abundance in the brain due to this limitation.

Another key advantage of MetaLigand is its automated pipeline for gene identification. By automatically identifying genes involved in the synthesis and transport of NPLs from comprehensive genomic and metabolomic databases, MetaLigand can collect a more exhaustive gene list than tools relying solely on manual curation. This automated approach improves the accuracy and scope of NPL detection, as seen in MetaLigand’s accurate identification of testosterone in steroidogenic tissues. Tools like CellphoneDBv5 and MRClinkDB missed important enzymes (e.g., HSD3B and CYP17A) and inaccurately predicted testosterone in non-steroidogenic tissues, such as the stomach.

## MATERIALS AND METHODS

### Data Collection Pipeline

A total of 3,088 unique metabolites were collected from HMDB (Version 5.0)^14^ using the “Browsing metabolites” page (https://hmdb.ca/metabolites) with the filtering terms “Endogenous” and “Detected and quantified”. All human receptors were first collected from 6 databases including OmniPath^24^, CellTalkDB^25^, Cellinker^20^, Cell-Cell Interaction Database^26^, NicheNet^5^, and NATMI-ConnectomeDB2020^27^ (Figure 1). After manual curation to filter out non-receptor proteins (e.g., calcium channels and metal ion transporters), 2,837 receptors were retained.

The collected human metabolites and receptors were then connected based on metabolite-protein interactions obtained from 6 databases: CTD^23^, BindingDB ^19^, Cellinker^20^, BioGRID^21^, STITCH^22^ and CellphoneDBv5^12^ (Figure 1). This resulted in 13,441 interaction pairs between 534 NPLs and 1,040 human receptors. Human receptor genes were then converted to mouse and zebrafish receptor genes based on orthologs using the ENSEMBL BioMart tool^64^.

To address the issue of varying metabolite nomenclature, NPLs were mapped to synonyms from the HMDB database to obtain HMDB accession IDs. HMDB accession IDs were used to query the genome-scale metabolic models (GEM) database^17^ for genes encoding synthetic and transport enzymes for each NPL.

Human, mouse, and zebrafish genes were collected from GEM-human, GEM-mouse, and GEM-zebrafish databases, respectively. Genes encoding synthetic enzymes or transporter proteins were collected from GEM model XML files using the following criteria: 1) Synthetic genes: filtered based on the reaction term “produce” and excluding the subsystem term “transport reactions”, 2) Efflux transporter genes: filtered by subsystem term “transport reactions” and compartment term “cytosol to extracellular”. The resulting gene list was then manually curated for accuracy. NPLs with shared synthetic genes (e.g., fatty acids, steroid hormones), genes for synthesizing NPL precursors (Synthetic step2) were included using the same filtering criteria. For NPLs with known salvage synthetic pathways (e.g., dopamine), genes encoding influx transporters for NPL precursors were incorporated by filtering for the subsystem term “transport reactions” and the compartment term “extracellular to cytosol”.

### NPL Predicting Model

The final genes collected for each NPL were categorized into five groups based on their functions: 1) Synthetic step 1: Genes encoding enzymes for synthesizing the NPL compound; 2) Synthetic step 2: Genes encoding enzymes for synthesizing the NPL precursors; 3) Efflux step: Genes encoding transporters for secreting the synthesized NPL from intracellular to extracellular location; 4) Influx step: Genes encoding proteins for transporting the NPL precursors from extracellular to intracellular location; 5) NPL receptors. Firstly, *De novo* synthesis pathway was estimated using the following formula:

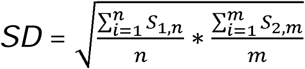

where S_1,n_ (n=1,2,3,4…) stands for list of genes encoding the enzymes for synthesizing the NPL compound (Synthetic step 1), S_2,m_ (m=1,2,3,4…) stands for list of genes encoding the enzymes for synthesizing the NPL precursor (Synthetic step 2). If a NPL do not have Synthetic step 2 genes, then the *de novo* synthesis pathway was simply estimated by:

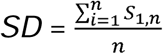

Then, salvage synthesis pathway was estimated using the following formula:

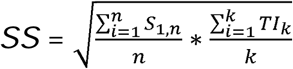

where *TI_k_* (k=1,2,3,4…) stands for list of genes encoding the transporters for transporting the NPL precursors from extracellular to intracellular location (Influx step). Then, the synthetic pathway of an NPL was then estimated by:

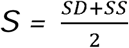

For NPLs without salvage synthesis pathway, their synthetic pathways were the same as *de novo* synthesis pathways (*S = SD*). Then, the final NPL abundance was estimated using the following formula:

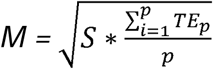

where *TE_p_* (p=1,2,3,4…) stands for list of genes encoding the transporters for transporting the NPL from intracellular to extracellular location (Efflux step). If an NPL do not have efflux transporters, its abundance was then estimated using only the synthetic pathway:

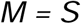

The cell-cell communication strength for each NPL-receptor interactions were determined using the following formula:

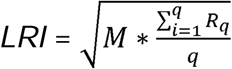

where M is predicted NPL abundance, and *R_q_* (q=1,2,3,4…) stands for list of genes encoding the receptors for an NPL.

## ACKNOWLEDGEMENT

This study was supported by National Eye Institute (NEI) grants R01EY031779 (JQ) and R01EY031685 (SB), and a Stein Innovation Award from Research to Prevent Blindness (SB).

## Figure legends

**Suppl. Fig.1.**
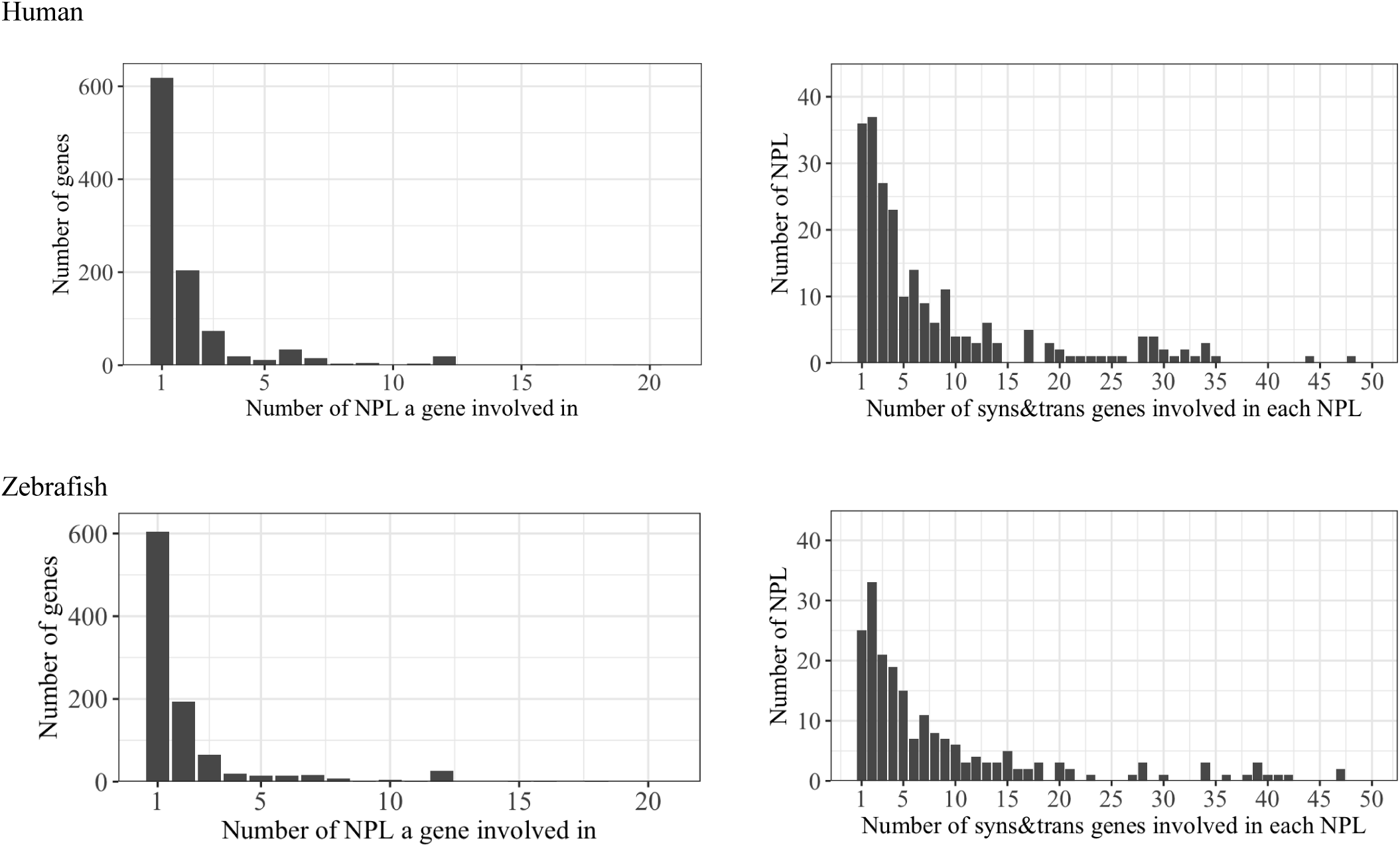
Histograms exhibit specificity of the association between NPLs and the relevant gene sets in MetaLigand human and zebrafish model. (Left) Distribution of the number of NPLs the genes involved in. (Right) Distribution of the number of synthetic enzymes and transporter genes involved in each NPL.

**Suppl. Fig.2.**
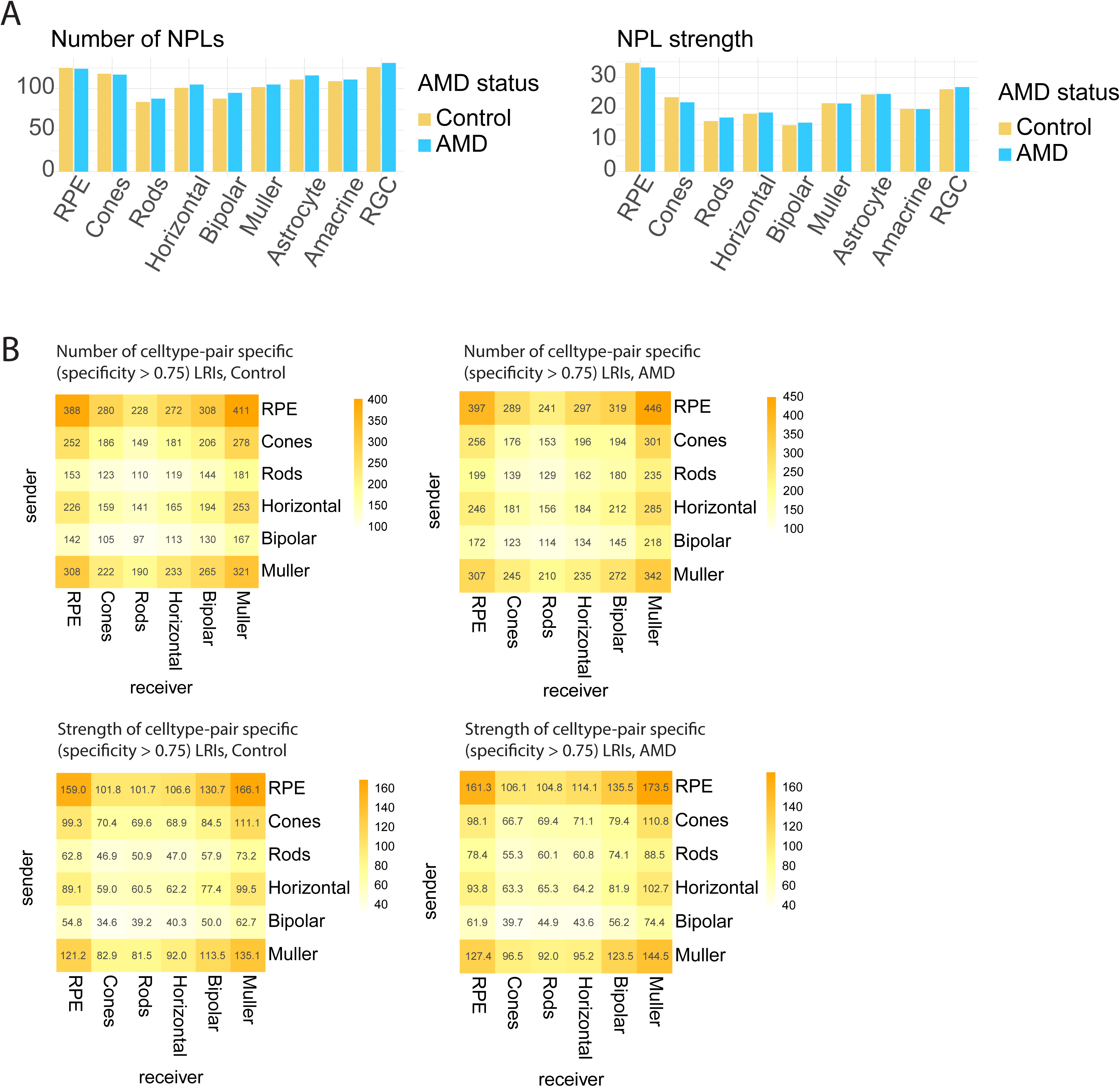
**(A)** Number (left) and strength (right) of NPLs (detected in at least 10% of the cells in the cluster of focus) predicted in each cell type in AMD and control cells. **(B)** Number (upper) and strength (lower) of NPL-mediated LRIs with specificity scores greater than 0.75 in control (left) and AMD (right) status between each pair of cell types.

**Suppl. Fig.3.**
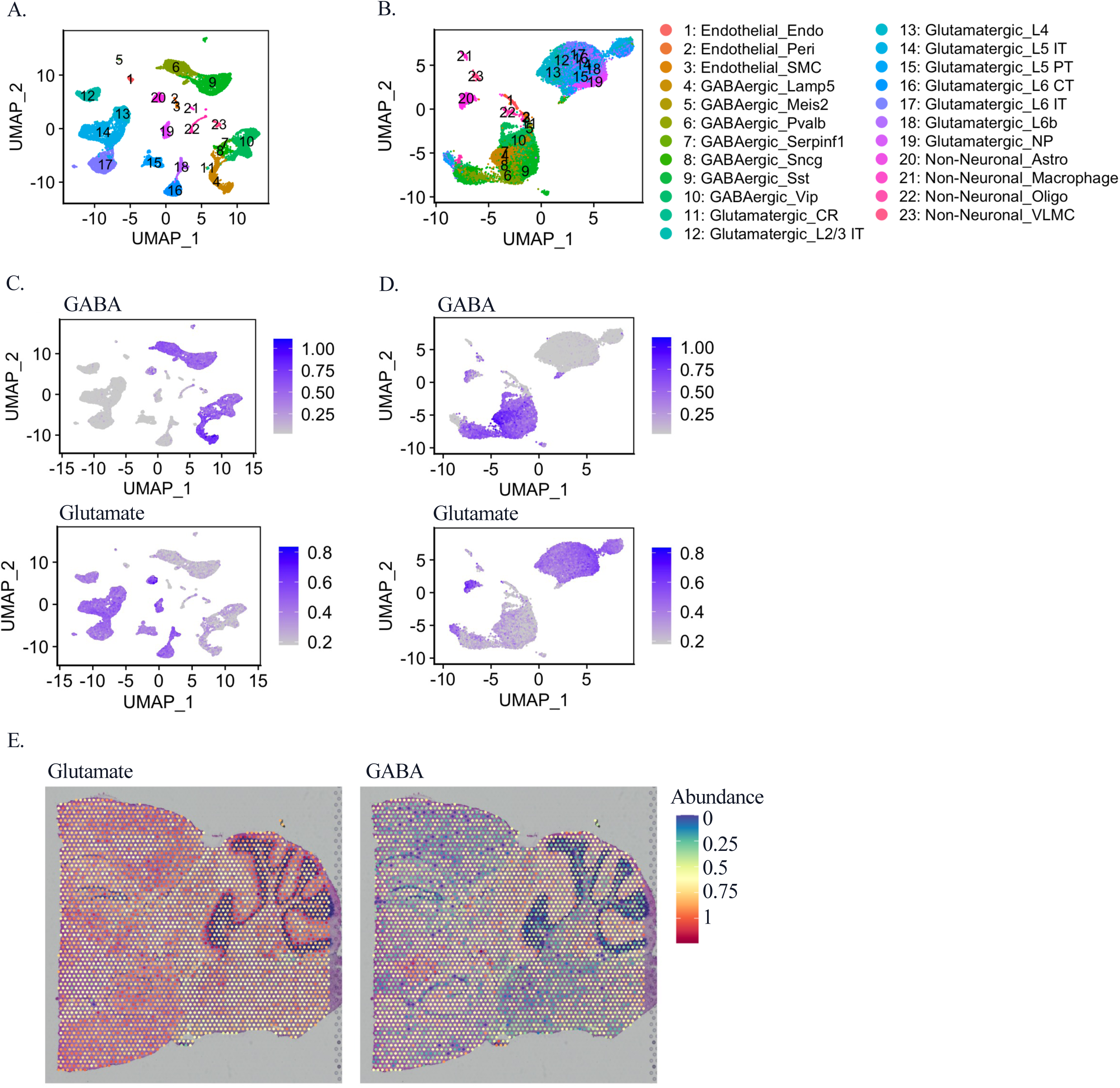
MetaLigand is compatible with popular computational tools for single-cell sequencing and spatial transcriptomics. **(A)** Cell clusters in the mouse brain Visp dataset visualized using UMAP dimensionality reduction by Seurat. **(B)** Cell clusters identified using MetaLigand predicted NPL levels as variable features for UMAP dimensionality reduction by Seurat. **(C-D)** Visualization of MetaLigand predicted GABA (upper) and glutamate (lower) levels and heterogeneity at single-cell resolution using UMAP by the FeaturePlot function of Seurat, with classical variable genes (left) or NPL levels (right) as variable features. **(E)** Scoring and projection of MetaLigand predicted glutamate (left) and GABA (right) levels onto slices from posterior brain segments using mouse brain slices data provided within the Seurat vignette.

